# DoAggregate: a pipeline for aggregation testing for rare-variant on a gene and pathway level

**DOI:** 10.64898/2026.10.29.762468

**Authors:** Simon Boutry, Cédric Van Marcke, Nisha Limaye, Guillaume Smits, Pascal Brouillard, Martina De Bortoli, Eleonore Pairet, Ali Saadat, Raphaël Helaers, Jacques Fellay, Tom Lenaerts, Miikka Vikkula

## Abstract

**Motivation:** Aggregation tests overcome the classical limitation of rare variant analysis in cohort study design. Choosing the right statistical test remains one of the biggest challenges. Existing tools fall short in offering a comprehensive solution that includes automatic data processing, testing across all genes, as well as exploration of broader genetic regions such as pathways and Gene Ontology (GO) terms. Most aggregation tests only report p-values and basic statistics, lacking in the presentation of cohort level statistics and comparison of results across whole genome or exome scales.

**Results:** We introduce DoAggregate, an innovative, scalable pipeline designed to address these challenges. DoAggregate enables users to perform aggregation tests across genes, pathways, and GO terms. It leverages Excalibur, an ensemble approach combining 36 distinct aggregation methods. The pipeline enhances data analysis by presenting over-representation analysis and aggregation results in detailed Excel sheets, complete with annotations and plots to facilitate nuanced interpretation and comparison. Applied to public exome data from the 1000 Genomes Project, DoAggregate tested 10,447 genes, 49 over-represented pathways and 36 over-represented GO terms in a single run, returning annotated, cohort-level results for each region.

**Availability and implementation:** DoAggregate source code, documentation and installation instructions as well as test data sets and results are freely available at https://github.com/boutrys/DoAggregate and released under the MIT license. The repository contains the serial implementation (R package), the parallel implementation for Slurm-based computer clusters and the web application.

**Supplementary information:** Supplementary data are available online.

## 1 Introduction

Over the last decade, methods for the association of rare variants, called aggregation tests, able to combine multiple markers within a genomic region have shown to be useful to overcome the limitations of classical GWASs (1–3). Despite their potential, the practical application of aggregation tests presents significant challenges, including data preprocessing, annotation, choosing the appropriate aggregation method, and the complex interpretation of results.

So far, several R packages integrating various aggregation methods have been proposed (*e.g*. SKAT, iGasso, WGScan, AssotesteR, RVTests) (4). In practice, these do not overcome the limitation of analysis preparation and results combinations when applied on a whole genome or exome. RAVAQ pipeline (5) is the first attempt to provide an integrated pipeline for rare variant association study. The main limitation of this pipeline is to only provide three aggregation tests and focus on only a single gene as the genetic region of interest. The evolution of these methods now permits analyses not just at the single-gene level but on groups of genes linked to biological pathways or shared GO terms (6,7), opening new avenues for understanding genetic contributions to disease. We introduce a comprehensive, user-friendly pipeline, DoAggregate, designed to facilitate the analysis of cohort data across genes, pathways, and GO terms. The user has access to 59 aggregation methods, in particular, the latest ensemble methods for aggregation testing (Excalibur) (8), DoAggregate provides interpretable results through detailed excel sheets and visual plots.

## 2 Materials and methods

Here we present DoAggregate, a pipeline (see Figure 1) for aggregation testing consisting of both a serial and parallelized implementation for parallel computing in a high-performance computing (HPC) environment, such as computer clusters. This pipeline uses 31 R packages as dependencies and the scheduler Slurm (9), when using the parallelized version. Using only two mandatory input parameters, the user can run the entire pipeline using one single command line (or R function). A web application, which lets users submit their cohort to the parallelized version through a browser and follow their jobs, is also provided. Useful resources such as R package installation scripts and publicly available datasets are provided in the GitHub repository (see Availability and implementation).

**Figure 1.**
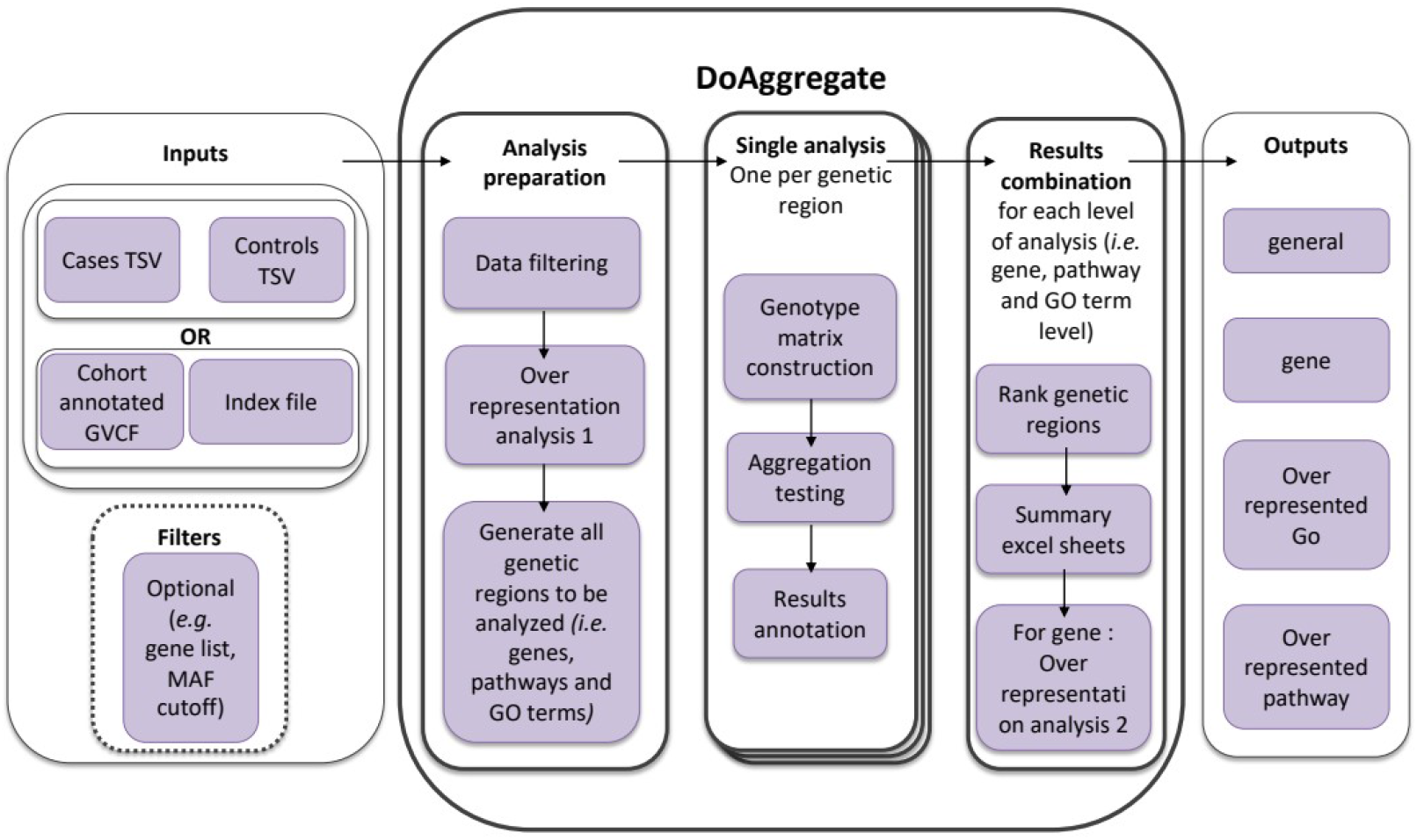
The figure presents the workflow of DoAggregate pipeline and data flow (black arrows). It outlines the process from input data through to the analysis and output stages, broken down into several key components. Inputs: The system accepts two main types of input data (Cases and controls data in TSV (Tab Separated Values) format or Cohort data in a Genome Variant Call Format (GVCF) file, along with an index file); Optional filters can be applied, such as specific gene lists or a minor allele frequency (MAF) cutoff. Analysis preparation: data filtering to refine the dataset; over-representation analysis of the filtered data; generation of a list of all genetic regions to be analyzed, which includes specific genes, pathways, and Gene Ontology (GO) terms. Single analysis: constructing a genotype matrix; performing aggregation testing; annotating the results. Results combination: ranking of genetic regions based on findings; creating summary excel sheets for detailed examination; further over-representation analysis specific to genes. Outputs: general output; gene-specific results; outputs identifying over-represented Gene Ontology (GO) terms and pathways, providing insights into biological functions and processes impacted by the genetic variants studied.

### 2.1 Inputs

Input data consists of annotated lists of genetic variants. This pipeline supports two input formats: (i) two tab separated (TSV) files (*i.e*. data on cases and controls), where each line matches a single variant per sample (*i.e*. a variant present in two samples will account for two lines in the input file); (ii) an annotated Genomic Variant Call Format (GVCF) file and an index file (tab separated text file) classifying individuals as cases or controls. Whatever the format, each input file must contain the following information for each variant: sample name or identifier, chromosome, chromosomal position, reference allele, alternative allele, gene symbol (annotated with HGNC database (10), also called SYMBOL when annotating using VEP (11)), zygosity (or genotype) and minor allele frequency (*e.g*. gnomad (12)). More advanced parameters can be used to refine the filtering (see Supplementary Data for mandatory columns, parameters and conversions).

### 2.2 Analysis preparation

The first task performed by DoAggregate is an optional data-cleaning, so only the relevant information is kept for the next steps (see detailed cleaning procedure in Supplementary Data). The default filtering applied only keeps rare variants with minor allele frequency less or equal to 0.01 that have existing gene annotation. Optional filters defined at launch will also be applied during this step. Based on all genes carrying at least one rare variant in the cohort (cases and controls), an over-representation analysis will be performed, based on the clusterProfiler R package (13,14) (see Supplementary Data for a detailed description of the over-representation analysis), exploring over-represented pathways from different databases (*i.e*. DiseaseOntology (15), DisGeNET (16), KEGG (17), NCG (18), Reactome (19) and WikiPathways (20)) and all GO terms (*i.e*. Biological process (BP), Cellular component (CC) and Molecular functions (MF)). At the end of this step, DoAggregate generates a list of all genetic regions (*i.e*. genes, significant over-represented pathways and GO terms) that will be tested for aggregation in the next step.

### 2.3 Single analysis

DoAggregate subsequently runs a single analysis per genetic region defined in the previous step. Input data needed for aggregation tests, such as a genotype matrix, will be generated. For aggregation testing, the default method is the Excalibur ensemble method, although the user has access to 59 specific aggregation tests. Each genetic region analyzed is annotated at the cohort level (*e.g*. number of mutations in cases and controls).

### 2.4 Results combination

As the previous step generates one set of results per genetic region, they need to be combined in an aggregated form. For this combination, DoAggregate defines three levels of analysis: (i) gene; (ii) over-represented pathway; (iii) over-represented GO term. The combination of all results at the same level of analysis comprises correction for multiple testing using Benjamini-Hochberg, ranking of genetic regions based on adjusted p-values, and identification of the specific statistical method that led to the identification of the genetic region. Significant genetic regions, in which the total number of variants in cases is equal or greater than the total of variant found in controls, are separated. For such genetic regions, an excel file containing the data of the original variants will be stored, allowing the user to directly investigate that region at the variant level. For gene level analyses, a second over representation analysis will be performed on the significant genes.

### 2.5 Outputs

DoAggregate outputs are separated into 4 folders (*i.e*. general, gene, over-represented pathway and over-represented GO, see Table S3 and supplementary). The general folder contains the cleaned data for case and control samples (if a cleaning was applied to input data) and the results of the first over-representation analysis. The three other folders will contain annotated Excalibur results for all genetic regions analyzed (in the form of Excel sheets) and a folder with the data behind the significant region (see Supplementary Data for a complete description of each file and annotations).

## 3 Results

### 3.1 Application to public exome data

To illustrate DoAggregate on real sequencing data, we ran the pipeline with default parameters on publicly available whole-exome data from the 1000 Genomes Project (23). We used the 26 individuals of the CEU population (Utah residents with Northern and Western European ancestry) as cases and the 25 individuals of the GBR population (British from England and Scotland) as controls. Both groups are of closely related European ancestry and carry no phenotype labels, so this comparison is close to a null setting: few significant associations are expected, and those found serve to illustrate the outputs rather than as biological findings. The joint-called VCF was annotated with ANNOVAR (24); genotypes with read depth below 10 were removed and variants with a gnomAD allele frequency of at most 0.01 (or absent from gnomAD) were kept (the scripts are provided with the test data).

After filtering, the dataset contained 81,242 unique rare variants (63,761 in cases and 60,184 in controls), corresponding to 210,949 and 196,058 variant calls in cases and controls, respectively. These variants fell in 12,969 genes, of which 10,447 (80.6%) carried at least two unique variants and were tested; the remaining 2,522 genes, each with a single variant, were reported as not analyzed. The over-representation analysis on the genes carrying at least one rare variant in the cohort retained 85 non-redundant significant terms (48 DisGeNET, 18 GO-BP, 9 GO-MF, 9 GO-CC, 1 WikiPathways), which defined 49 pathway regions and 36 GO-term regions for aggregation testing (Table 1, Figure 2a).

**Table 1.** Number of genetic regions defined, tested and found significant at each level of analysis for the 1000 Genomes exome example (26 CEU cases vs 25 GBR controls).

| **Level** | **Regions defined** | **Regions tested** | **Not tested (<2 variants)** | **Median tests applied** | **Excalibur q ≤ 0.05** | **of which more variants in cases** | **Best single-test q ≤ 0.05** | **Lowest Excalibur q** |
|---|---|---|---|---|---|---|---|---|
| Gene | 12969 | 10447 | 2522 | 30 | 5 | 3 | 37 | 0.000827 |
| Pathway | 49 | 49 | 0 | 27 | 0 | 0 | 5 | 0.318 |
| GO term | 36 | 36 | 0 | 27 | 2 | 1 | 7 | 0.0326 |

**Figure 2.**
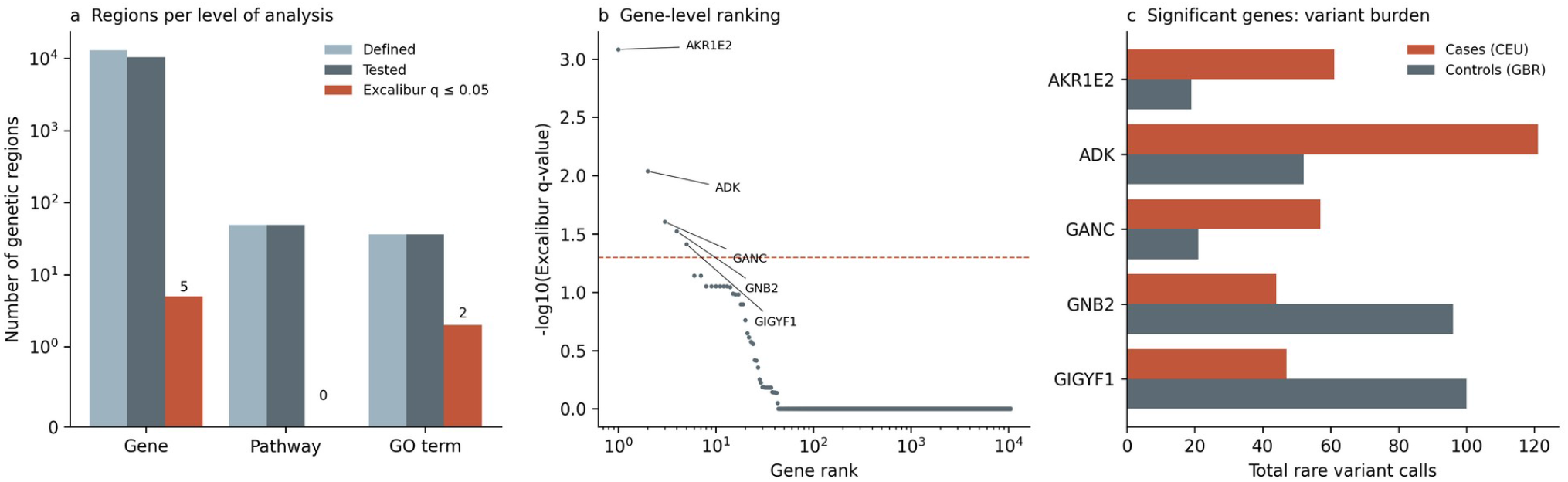
DoAggregate applied to 1000 Genomes exomes (26 CEU cases vs 25 GBR controls). (a) Number of genetic regions defined, tested and significant (Excalibur q ≤ 0.05) per level of analysis (symmetric log scale). (b) Genes ranked by Excalibur q-value; the dashed line marks q = 0.05. (c) Total number of rare variant calls in cases and controls for the five significant genes; only regions with at least as many variants in cases as in controls have their variant-level data exported.

At the gene level, 5 of 10,447 tested genes (0.05%) reached an Excalibur q-value of at most 0.05 (median q-value 1; Table 2, Figure 2b), consistent with the near-null design. For AKR1E2, ADK and GANC, the total number of variants was higher in cases than in controls, so DoAggregate exported their variant-level data for direct inspection; for GNB2 and GIGYF1, the signal was driven by an excess of variants in controls (Figure 2c). A median of 30 of the 36 Excalibur tests could be applied per gene (range 18–30), the others failing or not being applicable to the region. At the GO-term level, 2 of 36 regions were significant (endoplasmic reticulum tubular network, q = 0.033; low-density lipoprotein particle binding, q = 0.033), and no over-represented pathway reached significance (lowest q = 0.32).

**Table 2.** Genetic regions with an Excalibur q-value ≤ 0.05. “Best single-test q” is the lowest q-value obtained by any single aggregation test, and “Best test” is that test.

| **Level** | **Region** | **Excalibur q** | **Best single-test q** | **Best test** | **Tests applied** | **Cases with variant** | **Controls with variant** | **Variants in cases** | **Variants in controls** | **Unique variants** |
|---|---|---|---|---|---|---|---|---|---|---|
| Gene | AKR1E2 | 0.000827 | 3.06e-05 | RWAS | 28 | 12 | 14 | 61 | 19 | 21 |
| Gene | ADK | 0.00916 | 0.000763 | REBET | 25 | 17 | 16 | 121 | 52 | 24 |
| Gene | GANC | 0.0249 | 0.00154 | REBET | 24 | 17 | 12 | 57 | 21 | 13 |
| Gene | GNB2 | 0.03 | 0.00154 | REBET | 27 | 8 | 16 | 44 | 96 | 35 |
| Gene | GIGYF1 | 0.0386 | 0.00179 | REBET | 28 | 11 | 16 | 47 | 100 | 43 |
| GO term | endoplasmic reticulum tubular network | 0.0326 | 0.00126 | REBET | 27 | 26 | 21 | 297 | 215 | 113 |
| GO term | low-density lipoprotein particle binding | 0.0326 | 0.00126 | REBET | 27 | 20 | 24 | 143 | 196 | 76 |

Alongside the Excalibur q-value, DoAggregate reports the best q-value obtained by any single test. In this dataset, 37 genes had at least one individual test with q ≤ 0.05 (REBET for 29, RWAS for 8). Because this value is the minimum over about 30 tests, without correction for that selection, it is optimistic. It is reported to help interpretation, while prioritization should rely on the Excalibur q-value.

### 3.2 Test dataset and GVCF input

A reduced version of the same data, restricted to nine genes, is distributed with the package as a test dataset. It exercises the GVCF input route together with the coverage-based cleaning step. On this dataset, cleaning removed 3 poorly covered controls (22 to 19) and reduced the number of variant calls from 2,055 to 1,504, raising the reliability index from 0.35 to 0.42. Because the nine genes were chosen among the top-ranked genes of the full analysis, this dataset is meant to check the installation and the outputs, not to assess statistical performance.

## 4 Discussion

DoAggregate is an easy-to-use pipeline that integrates data processing, over-representation analyses, numerous aggregation tests (with the latest optimal combination of 36 aggregation tests, called Excalibur), and combination of results. Thanks to the available parallelized implementation, DoAggregate is a modern, scalable genetic region prioritization tool. In terms of computational time, we strongly recommend using the parallelized version, when possible, rather than the serial implementation, as the number of genetic regions tested grows with cohort size (e.g. more than 10,000 genes for the exome dataset in section 3).

This pipeline’s ability to process and analyze data at multiple levels – from individual genes to broader biological pathways and GO terms – enables a more nuanced understanding of the genetic architecture of diseases. It allows researchers to explore genetic data with unprecedented depth and precision. Furthermore, DoAggregate’s design addresses key challenges in the field, including the need for efficient data management, the ability to handle vast datasets, and the provision of clear, actionable results. By offering outputs that are both comprehensive and easy to interpret, it facilitates a deeper exploration of the biological significance of identified variants, potentially accelerating the discovery of biomarkers and therapeutic targets. We believe that combining DoAggregate with tools such as VarCoPP (21) and ORVAL (22) could speed up the discovery of digenic and oligogenic signatures.

We plan to employ three strategies to improve this pipeline in the future: integration of new aggregation tests and update of the ones already included, continuous improvement of the optimal ensemble Excalibur method, and improvement of data filtering by integrating RAVAQ strategy.

## Acknowledgements

We would like to thank all the members of the laboratory of Human Molecular Genetics and members of the oligogenic team at the Interuniversity Institute of Bioinformatics in Brussels for their support and feedback. We also thank the National Lottery, Belgium and the Foundation against Cancer (2010-101), Belgium for their support to the Genomics Platform of University of Louvain and de Duve Institute, as well as the Fonds de la Recherche Scientifique - FNRS Equipment Grant U.N035.17 for the «Big data analysis cluster for NGS at UCLouvain». S.B. was supported by fellowships from F.R.I.A. (Fonds pour la formation à la recherche dans l’industrie et dans l’agriculture), and Patrimoine UCL. The authors thank the Genomics Platform of University of Louvain for access to the biocomputing cluster.

## References

1. Uffelmann E, Huang QQ, Munung NS, de Vries J, Okada Y, Martin AR, et al. Genome-wide association studies. Vol. 1, Nature Reviews Methods Primers. Springer Nature; 2021.

2. Li B, Liu DJ, Leal SM. Identifying rare variants associated with complex traits via sequencing. Curr Protoc Hum Genet. 2013;(SUPPL.78).

3. Andrew R. Collins. Linkage Disequilibrium and Association Mapping. Methods Molecular Biololgy; 2007.

4. Boutry S, Helaers R, Lenaerts T, Vikkula M. Rare variant association on unrelated individuals in case-control studies using aggregation tests: existing methods and current limitations. Vol. 24, Briefings in Bioinformatics. Oxford University Press; 2023.

5. Marenne G, Ludwig TE, Bocher O, Herzig AF, Aloui C, Tournier-Lasserve E, et al. RAVAQ: An integrative pipeline from quality control to region-based rare variant association analysis. Genet Epidemiol. 2022 Jul 1;46(5–6):256–65.

6. Aleksander SA, Balhoff J, Carbon S, Cherry JM, Drabkin HJ, Ebert D, et al. The Gene Ontology knowledgebase in 2023. Genetics. 2023 May 1;224(1).

7. Ashburner M, Ball CA, Blake JA, Botstein D, Butler H, Cherry JM, et al. Gene ontology: Tool for the unification of biology. Vol. 25, Nature Genetics. 2000. p. 25–9.

8. Boutry S, Helaers R, Lenaerts T, Vikkula M. Excalibur: A new ensemble method based on an optimal combination of aggregation tests for rare-variant association testing for sequencing data. PLoS Comput Biol. 2023 Sep 1;19(9 September).

9. Jette M, Grondona M, Livermore L. Approved for public release; further dissemination unlimited SLURM: Simple Linux Utility for Resource Management [Internet]. 2003. Available from: http://www.llnl.gov/tid/Library.html

10. Seal RL, Braschi B, Gray K, Jones TEM, Tweedie S, Haim-Vilmovsky L, et al. Genenames.org: the HGNC resources in 2023. Nucleic Acids Res. 2023 Jan 6;51(D1):D1003–9.

11. McLaren W, Gil L, Hunt SE, Riat HS, Ritchie GRS, Thormann A, et al. The Ensembl Variant Effect Predictor. Genome Biol. 2016 Jun 6;17(1).

12. Chen S, Francioli LC, Goodrich JK, Collins RL, Kanai M, Wang Q, et al. A genomic mutational constraint map using variation in 76,156 human genomes. Nature. 2024 Jan 4;625(7993):92–100.

13. Wu T, Hu E, Xu S, Chen M, Guo P, Dai Z, et al. clusterProfiler 4.0: A universal enrichment tool for interpreting omics data. Innovation. 2021 Aug 28;2(3).

14. Yu G, Wang LG, Han Y, He QY. ClusterProfiler: An R package for comparing biological themes among gene clusters. OMICS. 2012 May 1;16(5):284–7.

15. Baron JA, Johnson CSB, Schor MA, Olley D, Nickel L, Felix V, et al. The DO-KB Knowledgebase: a 20-year journey developing the disease open science ecosystem. Nucleic Acids Res. 2024 Jan 5;52(D1):D1305–14.

16. Piñero J, Ramírez-Anguita JM, Saüch-Pitarch J, Ronzano F, Centeno E, Sanz F, et al. The DisGeNET knowledge platform for disease genomics: 2019 update. Nucleic Acids Res. 2020 Jan 1;48(D1):D845–55.

17. Kanehisa M, Furumichi M, Sato Y, Kawashima M, Ishiguro-Watanabe M. KEGG for taxonomy-based analysis of pathways and genomes. Nucleic Acids Res. 2023 Jan 6;51(D1):D587–92.

18. Dressler L, Bortolomeazzi M, Keddar MR, Misetic H, Sartini G, Acha-Sagredo A, et al. Comparative assessment of genes driving cancer and somatic evolution in non-cancer tissues: an update of the Network of Cancer Genes (NCG) resource. Genome Biol. 2022 Dec 1;23(1).

19. Milacic M, Beavers D, Conley P, Gong C, Gillespie M, Griss J, et al. The Reactome Pathway Knowledgebase 2024. Nucleic Acids Res. 2024 Jan 5;52(D1):D672–8.

20. Agrawal A, Balcı H, Hanspers K, Coort SL, Martens M, Slenter DN, et al. WikiPathways 2024: next generation pathway database. Nucleic Acids Res. 2024 Jan 5;52(D1):D679–89.

21. Versbraegen N, Gravel B, Nachtegael C, Renaux A, Verkinderen E, Nowé A, et al. Faster and more accurate pathogenic combination predictions with VarCoPP2.0. BMC Bioinformatics. 2023 Dec 1;24(1).

22. Renaux A, Papadimitriou S, Versbraegen N, Nachtegael C, Boutry S, Nowé A, et al. ORVAL: a novel platform for the prediction and exploration of disease-causing oligogenic variant combinations. Nucleic Acids Res. 2019 Jul 1;47(W1):W93–8.

23. 1000 Genomes Project Consortium, Auton A, Brooks LD, Durbin RM, Garrison EP, Kang HM, et al. A global reference for human genetic variation. Nature. 2015;526(7571):68–74.

24. Wang K, Li M, Hakonarson H. ANNOVAR: functional annotation of genetic variants from high-throughput sequencing data. Nucleic Acids Res. 2010;38(16):e164.

