## Supplementary Data for "DoAggregate: a pipeline for aggregation testing for rare-variant on a gene and pathway level"

### *S1 Input data*

DoAggregate accepts two input formats. Whatever the format, the information listed in Table S1 is required for every variant of every individual.

#### TSV files

Two tab-separated files are given, one for cases and one for controls, with one line per variant per individual (a variant carried by two individuals appears on two lines). Column names must match Table S1; any additional column is kept and reported in the variant-level output files.

#### Annotated VCF or GVCF file

A single annotated multi-sample VCF (or GVCF) is given together with two text files listing, one per line, the identifiers of the cases and of the controls. The argument mandatory_column_mapping (or options -0 to -9 of the cluster version) maps each mandatory field of Table S1 to the corresponding VCF field, for example gene_symbol = "Gene.ensGene" for an ANNOVAR annotation, "SYMBOL" for VEP, and gnomad_wes_af = "gnomad40_AF". The VCF is read with the vcfR package. When no zygosity field is given, zygosity is derived from the genotype (GT): 0/1 and 0|1 are heterozygous, 1/1 and 1|1 homozygous alternative, and 0/0 and 0|0 homozygous reference. Only heterozygous and homozygous alternative genotypes are kept as variants. The read depth (DP) of each genotype is used by the optional cleaning step (section S3).

#### Automatic conversions and checks

- Lines with a missing sample, chromosome, position, reference allele, alternative allele, gene symbol or zygosity are removed.
- Chromosomes coded 23 and 24 are renamed X and Y.
- When a variant is annotated to several genes (e.g. "GENE1\x3bGENE2" in ANNOVAR output), the first gene is kept.
- Variants with a population allele frequency above the rare-variant threshold (default 0.01) are removed. Variants absent from the population database (missing frequency) are kept.
- Variants are identified as chromosome_position_reference_alternative to build the genotype matrices.

#### Optional gene list

A text file with one HGNC gene symbol per line restricts the analysis to these genes.

**Table S1.** Mandatory information for each variant.

| Column | Description | Example |
|---|---|---|
| sample | Identifier of the individual | NA06984 |
| chr | Chromosome (with or without the chr prefix; 23/24 are converted to X/Y) | 1 |

| | | |
|---|---|---|
| pos | Chromosomal position | 721290 |
| reference | Reference allele | G |
| alternative | Alternative allele | C |
| zygosity | Heterozygous or Homozygous (TSV input); derived from GT for VCF input when not provided | Heterozygous |
| gene_symbol | HGNC gene symbol (SYMBOL in VEP, Gene.ensGene or Gene.refGene in ANNOVAR) | ADK |
| gnomad_wes_af | Population allele frequency of the alternative allele (e.g. gnomAD) | 0.0012 |
| gt_GT, gt_DP | VCF input only: genotype and read depth of each individual | 0/1, 35 |

### *S2 Parameters*

Table S2 lists the parameters of the serial R package (function main) and the corresponding options of the parallel version (submit_Excalibur.sh). Only the input files are mandatory; default values are used for all other parameters.

**Table S2.** Main parameters of DoAggregate.

| R argument | Cluster option | Default | Description |
|---|---|---|---|
| PATH_PATIENTS, PATH_CONTROLS | -A, -B | mandatory | Case and control TSV files, or lists of case and control identifiers for VCF input |
| PATH_GVCF | -G | none | Annotated VCF/GVCF file (VCF input only) |
| importation_method | -I | 0 | 0: two TSV files; 1: one VCF/GVCF file |
| mandatory_column_mapping | -0 to -9 | see S1 | Names of the VCF fields holding the mandatory information |
| path_results | last argument | none | Output folder |
| list_genes | -b | 0 (all genes) | File with a list of genes to analyse |
| rare_maf_threshold | -i | 0.01 | Maximum population allele frequency of a rare variant |
| min_var | -f | 2 | Minimum number of unique variants for a region to be tested |
| max_var | -g | 10,000 (serial), 100,000 (cluster) | Maximum number of unique variants for a region to be tested |

| | | | |
|---|---|---|---|
| remove_private | -x | 0 | 1: remove variants carried by a single individual |
| var_covariate_pca | -k | 1 | 1: use the first two principal components of the region genotype matrix as covariates |
| version | -u | "optimal" | "optimal": Excalibur and its 36 tests; "all": all 59 tests; or a list of test names |
| pvalue_threshold | | 0.05 | Significance threshold on Benjamini-Hochberg adjusted p-values (q-values) |
| DO_over_representation | | 1 | Run the over-representation analysis and test over-represented pathways and GO terms |
| do_data_preparation | -q | 1 | Run the coverage-based cleaning (section S3) |
| coverage_matrix_path | | 0 (none) | TSV input: coverage matrix with columns chr, pos and one read-depth column per individual |
| min_coverage | -n | 20 | Minimum read depth for a position to be well covered in an individual |
| mapqv_threshold | -m | 20 | Minimum mapping quality of reads counted when the coverage matrix is built from BAM files |
| remove_individual | -o | 1 | Remove badly covered individuals |
| remove_position | -p | 1 | Remove badly covered positions |
| (cluster only) | -E, -P, -Q, -R, -L, -M | | E-mail, maximum number of jobs, Slurm partition, R module, R library path and module command |

### *S3 Data cleaning procedure*

The optional cleaning step uses the sequencing depth of every variant position in every individual to remove individuals and positions that are poorly covered, because a missing variant call at such positions cannot be distinguished from a reference genotype. The depth is taken from the DP field of the VCF, or from a coverage matrix (positions by individuals) built from the BAM files counting only reads with a mapping quality of at least mapqv_threshold. The procedure is:

1. A position is considered badly covered in an individual when its depth is below min_coverage. The reliability index is the proportion of entries of the coverage matrix that reach min_coverage.

2. For each individual, the number of badly covered positions is counted (file ID 4, Figure S1a). Individuals for which this number exceeds the median plus one standard deviation over all individuals are removed (removed_cases.xlsx and removed_controls.xlsx), and the counts are recomputed (file ID 3).

3. For each position, the number of badly covered individuals is counted (file ID 1, Figure S1b). Positions for which this number exceeds the median plus one standard deviation over all positions are removed, and the counts are recomputed (files ID 2 and 5).

4. Variants that are no longer carried by any remaining individual are removed, and the reliability index is recomputed. File ID 6 (impact_of_cleaning.xlsx) reports the number of cases, controls, unique variants and variant calls, and the reliability index, before and after cleaning.

On the small public test dataset, cleaning removed 3 of 22 controls and reduced the number of variant calls from 2,055 to 1,504, raising the reliability index from 0.35 to 0.42.

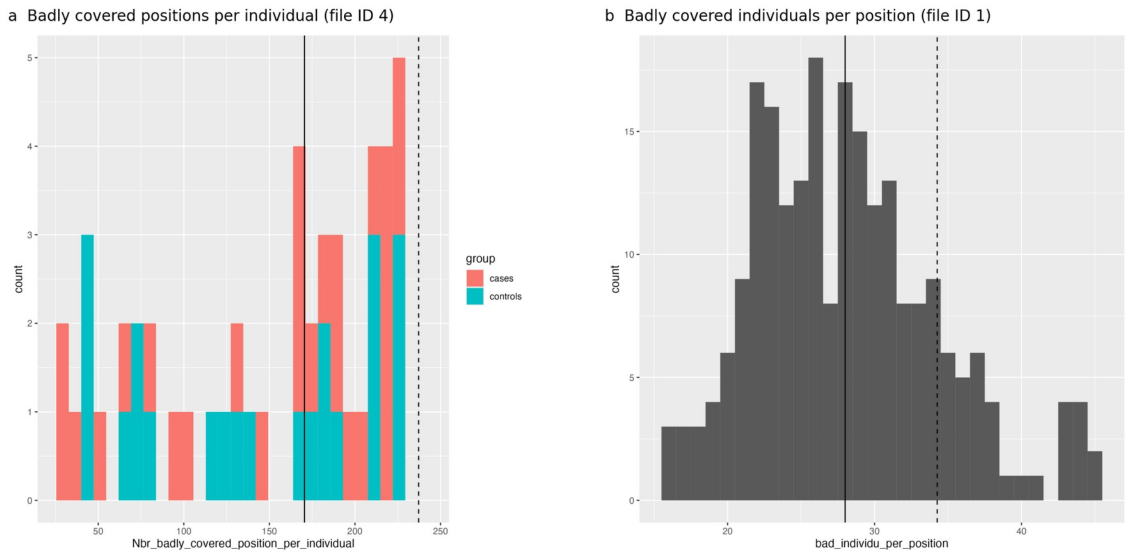


**Figure S1.** Coverage-based cleaning on the small public test dataset. (a) Number of badly covered positions (depth below min_coverage) per individual before removal, coloured by group. (b) Number of badly covered individuals per position. The solid and dashed lines mark the median and the median plus one standard deviation; individuals or positions beyond the dashed line are removed.

### *S4 Over-representation analysis*

The over-representation analysis is performed with the clusterProfiler R package on the genes that carry at least one rare variant after filtering and cleaning. Gene symbols are converted to Entrez identifiers with the bitr function and org.Hs.eg.db; genes that cannot be converted are listed in file ID 28. Over-representation is tested for the three GO ontologies (biological process, cellular component and molecular function; enrichGO), KEGG (enrichKEGG), Reactome (ReactomePA), Disease Ontology, the Network of Cancer Genes and DisGeNET (DOSE), and WikiPathways. The current cluster version instead tests the gene sets of the MSigDB collections (tables in cluster/Data) with the enricher function and applies the same filters. P-values are adjusted with the Benjamini-Hochberg procedure, and terms with an adjusted p-value of at most 0.05 are kept.

Results are then simplified: (i) terms containing more input genes than max_gene_per_item (20 in the serial package, 500 in the cluster version) are removed to avoid very broad terms, as are terms containing a single input gene; (ii) for GO terms, the semantic similarity between terms is computed with GOSemSim and the term with the highest similarity to the others is removed recursively until no pair exceeds 0.5; (iii) terms mapping exactly the same list of input genes are collapsed into one line, keeping the most significant term. All significant terms of all databases are finally combined in file ID 34, and each resulting gene list is tested as one genetic region at the over-represented pathway or GO-term level. The output files are described in section S6.

### *S5 Aggregation testing and combination of results*

For each genetic region (gene, over-represented pathway or GO term), DoAggregate builds the genotype matrix of all cases and controls (0, 1 or 2 copies of the alternative allele for each rare variant of the region) and a binary phenotype (1 for cases, 0 for controls). Regions with fewer than min_var or more than max_var unique variants are not tested and are listed in general_info_not_analyzed. Population allele frequencies are used as variant weights by the weighted tests, and by default the first two principal components of the region genotype matrix are used as covariates.

With the default version, the 36 aggregation tests that compose Excalibur are applied: burden, SKAT and SKAT-O with robust p-values (SKAT package); ASCORE, ASSU and ASUM and their ordered versions, BST, C-alpha, CAST, CMAT, CMC, RBT, RVT1, RVT2, RWAS, SCORE, SSU, SSUW, the asymptotic SCORE, SSU, SSUW and t-tests, VT, WSS and WST (AssotesteR); KAT; REBET; and the RR, SPLS, T1P, T5P, WEP and SCORE-VTP tests. Tests that fail or are not applicable to a region return no p-value, and the number of tests successfully applied is reported for each region. Within a region, the p-values of the applied tests are adjusted with the Bonferroni correction, and the Excalibur p-value is the smallest adjusted value.

At each level of analysis, the p-values of each test, including Excalibur, are then adjusted across all tested regions with the Benjamini-Hochberg procedure. Regions are ranked by their Excalibur q-value, and those with a q-value of at most pvalue_threshold are reported as significant. For each region, DoAggregate also reports the lowest q-value obtained by any single test, that test, and the list of tests with a significant q-value. Because the lowest single-test q-value is selected among about 30 tests, it should be used for interpretation only, and prioritization should rely on the Excalibur q-value.

### *S6 Results description*

In this section we describe the results folders and files that DoAggregate will return as output. For this section, the name of folders will be in bold (*e.g.* **NameOfFolder**) and in italic (*e.g. exampleOfFile*) if this file is optional (*i.e.* it might not be present in some analysis results). For example, if no genes with significant aggregation were found, then the file *significant general info* will not exist in the folder **Gene**.

| File ID | Folder name | Sub folder or file name | Sub folder or file | File name |
|---|---|---|---|---|
| 1 | | *bad_individual_per_position.png* | | |
| 2 | | *bad_individual_per_position_after_position_removal.png* | | |
| 3 | | *bad_pos_per_individuals_after_individu_removal.png* | | |
| 4 | | *bad_pos_per_individuals_before_removal.png* | | |
| 5 | | *bad_pos_per_individuals_final.png* | | |
| 6 | | *impact_of_cleaning.xlsx* | | |
| 7 | | | | *BP_GO_over_representation.xlsx* |
| 8 | | | | *BP_GO_removed_duplicated_gene_list.xlsx* |
| 9 | | | | *BP_GO_removed_similar_over_representation.xlsx* |
| 10 | | | **BP** | *BP_GO_removed_too_big.xlsx* |
| 11 | | **GO_over_representation** | | *Dot_plot_BP_GO_overrepresentation.png* |
| 12 | | | | *GO_plot_BP_GO_overrepresentation.png* |
| 13 | | | | *Plot_Gene_occurence_GO_BP.png* |
| 14 | | | **CC** | *same files as file ID 7 to 13* |
| 15 | | | **MF** | *same files as file ID 7 to 13* |
| 16 | | | | *same file as file ID 7* |
| 17 | | | | *same file as file ID 8* |
| 18 | **General** | | | *same file as file ID 10* |
| 19 | | | **DiseaseOntology** | *same file as file ID 11* |
| 20 | | | | *Network_Circle_DiseaseOntology_overrepresentation.png* |
| 21 | | **pathway_over_representation** | | *Network_DiseaseOntology_overrepresentation.png* |
| 22 | | | | *same file as file ID 13* |
| 23 | | | **DisGeNET** | *same files as file ID 16 to 22* |
| 24 | | | **KEGG** | *same files as file ID 16 to 22* |
| 25 | | | **NetCanGen** | *same files as file ID 16 to 22* |
| 26 | | | **Reactome** | *same files as file ID 16 to 22* |
| 27 | | | **WikiPathways** | *same files as file ID 16 to 22* |
| 28 | | *list_gene_not_handled_for_analysis.xlsx* | | |
| 29 | | *Combined_removed_duplicated_gene_list.xlsx* | | |
| 30 | | *Gene_not_over_represented.xlsx* | | |
| 31 | | *Dot_plot_combined_overrepresentation.png* | | |
| 32 | | *Cooccurence_genes.xlsx* | | |
| 33 | | *Cooccurence_genes.png* | | |
| 34 | | *Summary_results_over_representation.xlsx* | | |
| 35 | | global_info_analysis.xlsx | | |
| 36 | | *general_info_not_analyzed.xlsx* | | |
| 37 | | *general_info.xlsx* | | |
| 38 | | *significant_general_info.xlsx* | | |
| 39 | **Gene** | *Cases_significant_results.xlsx* | | |
| 40 | | ***Sig_gene_over_representation*** | *produce same folder and files as file ID 7 to 34* | |
| 41 | | ***Cases_Sig_gene_over_representation*** | *produce same folder and files as file ID 7 to 34* | |
| 42 | | ***data_significant_region*** | NameCaseSigGeneticRegion.xslx | |
| 43 | **over_represented_pathway** | *produce same files as file ID 36 to 39 and folder ID 42* | | |
| 44 | **over_represented_GO** | *produce same files as file ID 36 to 39 and folder ID 42* | | |

**Table S3**. Schema of the results folders and files from DoAggregate. In italic, folder/file that might not be created. In bold, name of folder.

General

This folder contains the results from the cleaning process, the over representation analysis and global information about the three levels of analysis (*i.e.* gene, overrepresented pathway and overrepresented GO terms, and the three other results folders).

File IDs 1 to 6 (see Table S3) correspond to the cleaning process using the coverage matrix. File IDs 3 to 5 are histograms showing the number of positions badly covered per individual, after removing individuals, before, and after removing badly covered position, respectively. File ID 4 is first to be produced. File ID 1 is a histogram (see Supplementary Figure S1) showing the number of badly covered individuals per position. While file ID 2 gives the same after having removed positions above the threshold (*i.e.* above median of badly covered individuals per position, plus on standard deviation). File ID 6 is a table giving the number of patients, controls, unique variants (each variant is counted once, no matter if several individuals have the variant), total variants (in this case each variant per individual is counted) before and after cleaning. It also gives a reliability index computed as the number of entries in the coverage matrix for which the minimum coverage and mapping quality are respected, divided by the number of total entries in that matrix. In other words, it corresponds to the percentage of data that meets the quality standards.

Next we describe all the results produced by the over-representation analysis. The first step to use clusterProfiler is to convert gene identifiers using bitr function. When an input gene (with HGNC annotation) cannot be mapped to an ENTREZID (1), DoAggregate stores these genes to file ID 28. Files ID 7 to 13 are generated by the over-representation analysis for the GO term on the BP ontology using clusterProfiler R package. From these results, GO terms that exceed the maximum number of genes from the input data for that term are first removed (see file ID 10) (Using the Count from clusterProfiler results and our input parameter max_gene_per_item, by default 20 in order to avoid broad pathways and GO terms). Then, using GOSemSim R package, DoAggregate computes similarity in between the remaining GO terms and remove recursively GO terms with the highest similarity with others, until the maximum similarity falls below 50% (default input parameter value) to obtain file ID 9. DoAggregate keeps GO terms with adjusted p-values below or equal to 0.05 and collapse (using an underscore) into a single term the id and description of term corresponding to the same gene list (see file ID 7).

All terms that have been collapsed using the procedure can be found in file ID 8. Using clusterProfiler plot functions, plots of the top 20 (default input parameter value) most significant GO terms (see file ID 11, 12 and 13) are generated. The **CC** and **MF** folders follow the same strategy (see file ID 14 and 15, respectively). Only files ID 9 and 12 are specifique to GO term over representation, all others will be found in the results of each database for which DoAggregate tests for over-representation analysis (see file ID 16 to 19 and 22). Only two plots (from clusterProfiler R package) are specific to pathways and display the top 20 most significant over represented terms (see file ID 20 and 21). DoAggregate uses the same strategy for all pathway databases (see file ID 23 to 27). Subsequently, DoAggregate combines all significant results from all GO term ontologies and pathway databases (all files from file ID 7 to 27 similar to file ID 7) and remove the least significant (based on the adjusted p-value) pathways/GO terms mapping to the same gene list (see file ID 29). DoAggregate collapses (using an underscore) all the removed IDs, descriptions and databases into a single line, adding the columns giving the information of how many IDs and databases mapped to the gene list (see file ID 34). The top 20 terms are plotted (default input parameters), generating file ID 31. From our input gene list for the over representation analysis, removing all genes that were not mapped (see file ID 28), all genes that are never found in any significantly overrepresented pathways/GO terms (see file ID 34) are stored to generate file ID 30. Finally, using the top 100 (default value parameter, to be used carefully as it can lead to explosion computational time) significant results found in file ID 34, DoAggregate builds a co-occurrence matrix that indicates on the diagonal the number of time out of 100 that a gene occurs in overrepresented pathways/GO terms.

The other part of the matrix shows how many times out of 100, two genes co-occur. These results can be found in file ID 32 and are plotted in file ID 33 using Complex Heatmap R package (2,3). In file ID 34, all gene lists that can be mapped to a GO term are used as overrepresented regions to be tested for aggregation and results for such genetic regions can be found in folder ID 44. All other gene lists (that only map to an overrepresented pathway in file ID 34) are tested for aggregation and results are found in folder 43.

For each level of analysis (*i.e.* gene, overrepresented pathway, and overrepresented GO terms), global information is stored in file ID 35 including:

- Number of genetic regions: at the start of analysis; analyzed; significant; and not analyzed.
- Number of unique positions (defined as chromosome and position): total; in patients; in controls

- Number of unique variants (defined as chromosome, position, reference and alternative): total; in patients; in controls
- Total number of variants in patients; and in controls

Gene

File ID 36 contains all genetic regions that were discarded from the analysis. By default, if there are less than 2 unique variants (no aggregation possible) or more than 100 000 unique variants (computational time explosion), a region is not analyzed. This file introduced a set of annotations for each genetic region:

- Number of tests successfully applied (remember that Excalibur is made of 36 aggregation methods).
- Name of the genetic region
- Number of patients and controls with at least one variant
- Proportion of patients and controls with at least one variant
- Total number of variants within the cohort, in the patients, and in the controls
- Number of unique variants (chromosome, position, reference, alternative) in the cohort, in the patients, in the controls, and found in both
- Total number of heterozygous and homozygous variants in patients, and in controls
- Proportion of reference, heterozygous, and homozygous variants in the cohort

All these annotations are also found in file ID 37 to 39. These files introduce additional annotations, such as:

- Ranking : out of all genetic regions analyzed, a ranking based on Excalibur q-value (p-value adjusted using Benjamini-Hochberg) is assigned
- Excalibur q-value
- Best q-value (out of all aggregation methods applied)
- The aggregation test that achieved the best q-value
- List of all aggregation tests that where applied and had a significant p-value
- Number of aggregation tests that returned a significant p-value

File ID 37 contains all the genetic regions that were analyzed, while file ID 38 contains only the regions with an Excalibur q-value less or equal to 0.05. Based on file ID 38, file ID 39 keeps the results for which the total number of variants in patients is equal or greater than the total number of variants in controls.

Taking all genes from file ID 38, DoAggregate performs an overrepresentation test (same as above, see files ID 7 to 34) in order to see whether genes with significant aggregation are biologically linked in some pathways or GO terms (see folder ID 40). Folder ID 41 follows the same logic, but focuses on significant genes where total number of variants is equal or higher in patients than in controls.

Based on file ID 39, all input data (variants and annotations as they were given as input by the user) regarding the significant genetic regions will be stored in a separate file within folder ID 42. For example, if GeneA and GeneB were the only two genetic regions left in file ID 39, then within folder ID 42, two excel files, called GeneA.xlsx and GeneB.xlsx, will be stored, filled with all the cohort data that was used to test these regions. This allows the user to directly explore the variant level data.

Overrepresented Pathways and Overrepresented GO Terms

These two folders will take as input (genetic regions to be analyzed) all the overrepresented pathways and GO terms identified in file ID 34, and gives the same files and folders as described above (see files ID 36 to 39, and folder ID 42).
